# A 3D Tumor-on-a-chip Platform to Identify Drugs that Block Breast Cancer Cell Intravasation

**DOI:** 10.64898/2026.03.19.712923

**Authors:** Narmada Perera, Diogo Coutinho, Carolina Morais, Maria Faria, Raquel Neto, William Roman, Edgar R. Gomes, Cláudio A. Franco, Luís Costa, David Barata, Karine Serre, Sérgio Dias, Ana Magalhães

## Abstract

Metastasis is the leading cause of death in breast cancer patients, yet there are no drugs specifically designed to block cancer cell intravasation, an early step of the metastatic cascade that originates circulating tumour cells (CTCs). A major challenge in developing anti-intravasation drugs is the scarcity of relevant in vitro platforms suitable for predictable drug discovery. Intravasation is a fundamental step of metastasis and involves the crossing of cancer cells through an endothelial barrier to enter the blood circulation. Here we developed an intravasation-on-a-chip model with controlled extracellular matrix composition, fluid flow and shear stress, which mimics the dynamic tumour-endothelium interface. The systems allows real-time imaging of intravasation and the isolation and quantification of intravasated cancer cells. As a proof-of-concept for drug testing, we show that perfusion with the PI3K/mTOR inhibitor Dactolisib, significantly reduced intravasation without compromising endothelial cell viability. The system also provides the capability to evaluate inhibitor on-target activity via imaging analysis. This intravasation-on-a-chip model offers a powerful, scalable, and imaging-compatible platform for discovering and evaluating anti-intravasation compounds.

## Introduction

Metastasis accounts for most breast cancer deaths, making strategies to prevent metastatic spread an urgent clinical priority^1,2^. Two initial steps of the metastatic cascade are the invasion of the surrounding extracellular matrix (ECM), and the crossing of blood vessel endothelial barrier (intravasation)^3^. This results in the release of cancer cells into circulation, which later exit the blood and colonize distant organs ^4,5^. Intravasation occurs not only from primary tumors, but also from established metastases which can originate additional metastases or even re-seed the primary sites^6,7^. Moreover, apart from originating new metastases, circulating tumor cells (CTCs) are also in the genesis of other pathological events such as the formation of blood clots^8,9^. A recent study that analyzed autopsies of cancer patients, revealed a high percentage of blood clots, as well as a peak of CTC release few weeks before patient death^10,11^. Therefore, blocking intravasation could prevent metastasis and the overall pathological consequences of having cancer cells in circulation.

Unfortunately, there is a deficit of strategies aimed at preventing intravasation of cancer cells, resulting from the difficulty posed by studying this process mechanistically. There are multiple factors that may contribute to cancer cell invasion and intravasation, namely the molecular and biophysical composition of the ECM, the action of other cell types within the tumor microenvironment (particularly endothelial cells), and the level of blood flow shear stress on the tumor microvasculature^12,13^.

Intravasation is generally assessed *in vivo*, either using intravital imaging of tumors in living animals and/or the quantification of circulating tumor cells ^4,12^. Both approaches, however, offer limited experimental control, making it difficult to systematically isolate individual factors or test multiple therapeutic compounds. *In vitro*, a major challenge in studying intravasation is the lack of suitable models that recapitulate tumor microphysiology. Traditional 2D cell culture systems poorly replicate this process due to lack of complexity, while 3D models such as organoids (featuring multicellular constructs and ECM-analogs) lack key features of intravasation, like perfusable vasculature, capabilities to visualize in real-time the intravasation process at single-cell level and have limited capacity for isolating the intravasated cells.

Recent technological advancements have led to the development of microphysiological systems that address many of the limitations associated with traditional 2D and 3D cell culture models. These systems incorporate compartmentalized but interconnected perfusable microchannels that allow precise control over multiple individual variables present and better replicate the scale of *in vivo* conditions ^14,15^. As such, factors such as cell and matrix composition, stiffness, fluid dynamics, shear stress, nutrient and biochemical gradients, can be modulated according to the hypothesis to be tested. Using this approach, 3D tumor-on-a-chip and vessel-on-a-chip models that mimic the dynamic interactions between tumor cells and blood vessels during intravasation can be generated ^16–20^. Here, we present an intravasation-on-a-chip model that can be used as an *in vitro* testing platform for anti-intravasation compounds. The system includes microscopic analysis allowing built-in capacity of on-target efficacy of the compound. Moreover, the presence of endothelial cells enables simultaneous safety assessment by monitoring vascular toxicity.

## Results

### Mimicking ECM directional invasion and intravasation using a tumor-on-a-chip model

To investigate the process of breast cancer cell invasion and intravasation, we bioengineered a tumor cell-blood vessel interface. This approach aimed at embedding metastatic breast cancer cells on an ECM-analogue, i.e. Matrigel, on the proximity of a perfusable microfluidic channel where endothelial cells recreate a lumen forming a vessel-like barrier. The system was mounted on top of glass to allow high-resolution imaging. The device is made of polydimethylsiloxane (PDMS) and composed of five parallel and independent channels, separated by PDMS trapezoid pillars, used to hold the hydrogel prior to crosslinking. The distance between the pillars allows the compartmentalization of different cell types while at the same time enabling their interaction (Figures 1A and 1B). The distal position of the cell types in the device was optimized to ensure cancer cell-directed migration and invasion through the ECM towards a nutrient source, i.e. the vessel. As such, we positioned green fluorescent protein (GFP) expressing metastatic breast cancer cells (MDA-MB-231) embedded in Matrigel in the central chamber of the chip, flanked by two Matrigel-filled channels. The last two channels were then filled with either Matrigel (first channel) or complete tumor cell media (fifth channel) (Figure 2A). This asymmetric configuration showed that cancer cells preferentially migrated directionally toward the serum-containing complete media channel (Figure 2B). This directional behavior is consistent with the establishment of a chemotactic gradient that promotes tumor cell motility within the three-dimensional extracellular matrix. No migration was observed when the non-metastatic breast cancer cell line, MCF-7, was seeded in the same conditions (Figure 2C), demonstrating the usefulness of the interface when comparing different cancer cell types or chemotactic factors.

**FIGURE 1.**
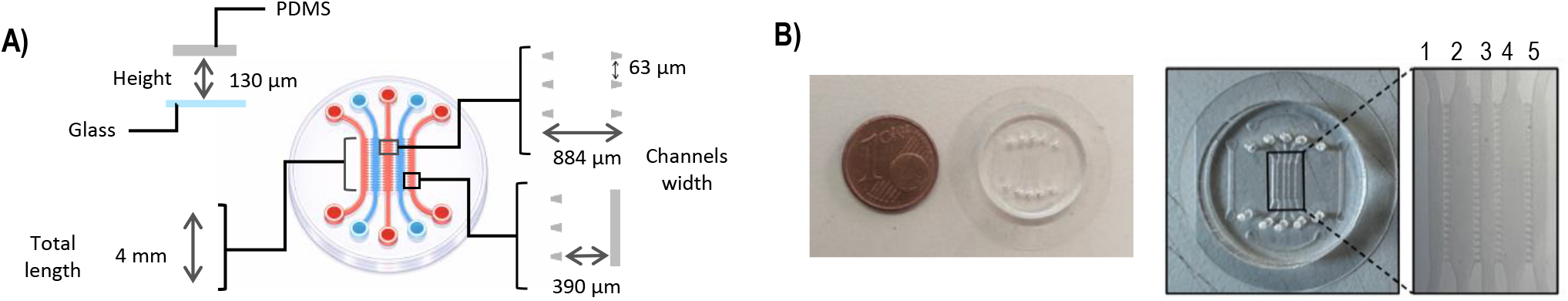
Characteristics of the device. **A)** Detailed dimensional specifications of the five-channel microfluidic device (created with BioRender.com). **B)** Images of the device, showing the five perfusable and interconnected microchannels.

**FIGURE 2.**
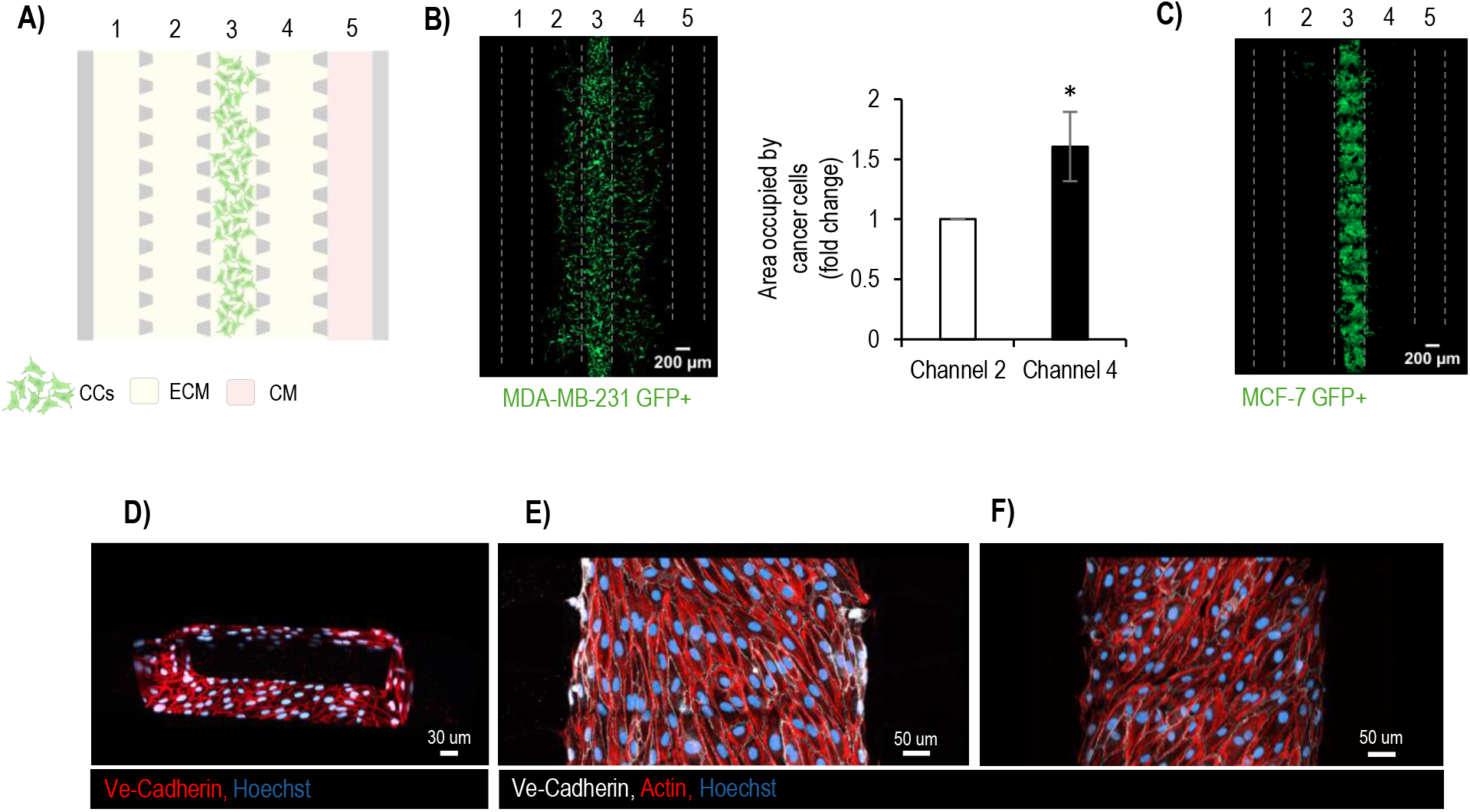
Establishment of the model. **A)** Schematic illustration of the tumor-on-a-chip platform, highlighting the configuration of each microfluidic channel on day 1 (created with BioRender.com). Channels 1, 2 and 4 contain Matrigel (ECM), channel 3 is seeded with cancer cells (CCs) embeded on Matrigel and channel 5 is filled with cancer cell complete media (CM). **B)** All-chip microscopic image captured on day three post-seeding, showing MDA-MB-231 GFP^+^ breast cancer cells (green) within the microfluidic device. Quantification of the area occupied by cancer cells on channels number 2 and number 4. The graph shows the median ± standard deviation of four chips and statistical significance was assessed by the two-sided t-test (* p<0.05). Scale bar = 200µm **C)** Representative image of MCF-7 GFP+ cells on the chip, three days after seeding. Scale bar = 200µm **D)** 3D reconstruction of confocal images showing endothelial cells (HUVECS) seeded on the chip. Confocal images of front view stained for VE-cadherin (red) and Hoechst (blue). Scale bar = 30µm. **E) and F)** bottom and top surfaces of the chip showing endothelial cells stained for actin (red) and VE-cadherin (white) and Hoechst (blue). Scale bar = 50µm.

To establish the microvessel, we seeded endothelial cells into the fifth channel. To validate vessel integrity, a 3D reconstruction of confocal Z-stacks demonstrated that endothelial cells formed a monolayer lining all four sides of the central chamber, as indicated by positive VE-cadherin staining (Figure 2D). High-resolution confocal planes of the top and bottom surfaces showed endothelial organization with co-staining of VE-cadherin and actin, confirming cytoskeletal integrity and cell-cell junction formation (Figure 2E and F). Taken together, we established and described the structural and functional assembly of a novel microfluidic tumor-vascular interface.

### Real-time imaging of intravasation

To visualize the dynamic interactions between metastatic breast cancer cells and the vascular endothelium during intravasation, we performed time-lapse confocal microscopy followed by 3D reconstruction of Z-stack image data (Figure 3). We were able to capture GFP-labeled cancer cells while they engage and cross the endothelial wall.

**FIGURE 3.**
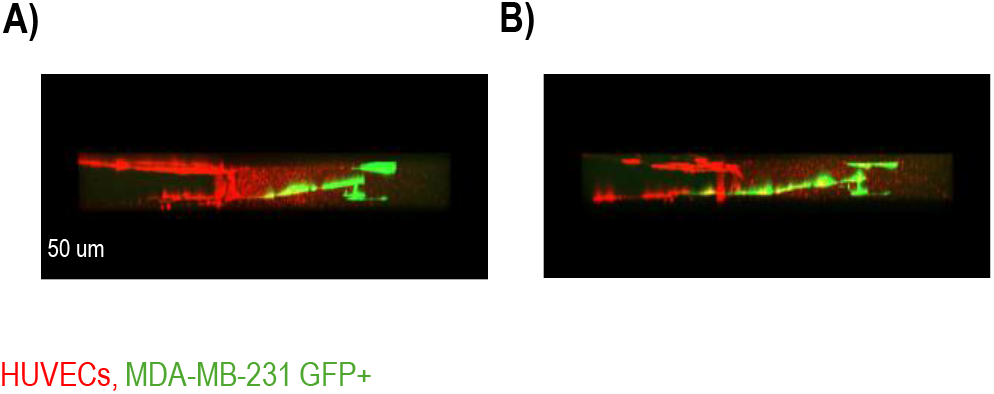
Time lapse imaging of tumor cell-endothelial interaction and intravasation. Front views of MDA-MB-231 GFP^+^ breast cancer cells (green) interacting with the endothelial cells (HUVECs-red). At 0 hours, one cancer cell establishes a contact with an endothelial cell (A), while at 10 hours it crossed the endothelial barrier (B). Images represent a 3D reconstruction of Z-stack confocal microscopy data. Scale bar = 50µm.

### Quantification and isolation of intravasated cells upon perfusion

Subsequently, we included perfusion of the vessel, which allowed the quantification and isolation of intravasated breast cancer cells in the collected effluent. The flow rate was increased gradually, as follows: 1□L/minute for one hour, followed by 2□L/minute for two hours, 3□L/minute for three hours and finally 6□L/minute for 18 hours. At the end of the experiment, cells on the chip were fixed by perfusing the endothelial chamber with paraformaldehyde and then stained and imaged (Figure 4A). Imaging analysis revealed a significantly greater number of GFP-positive cancer cells at the outlets compared to the inlets of the chip (Figure 4B and C). Moreover, we successfully isolated intravasated GFP-positive cancer cells at the end of the perfusion period (Figure 4B). These intravasated cells were alive and proliferative (not shown). Quantification of the number of cancer cells in the outlet, plus the collected cancer cells, led to the definition of an intravasation score. This allowed using our system to quantitatively test the anti-intravasation efficacy of model drugs.

**FIGURE 4.**
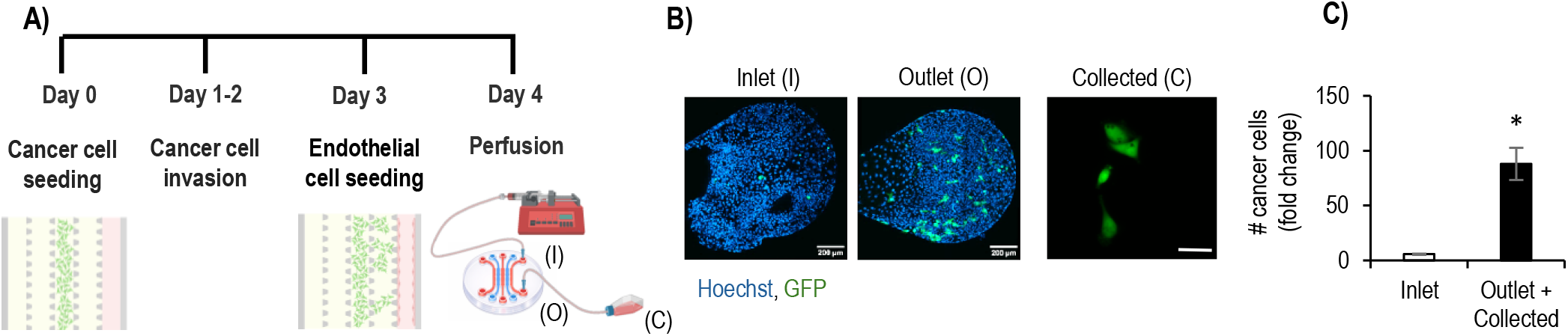
Perfusion and quantification of intravasation. **A)** Schematic representation of the experimental steps of the assay and perfusion system (created with BioRender.com). **B)** Representative images of an experiment showing inlets (I), outlets (O) and collected cells (C). **C)** Quantification of cancer cells at the three locations, Scale bar = 200 μm in I and O, and 50 μm in (C). The graph presents the mean ± standard deviation from three independent experiments and statistical significance was assessed by the two-sided t-test (* p<0.05).

### Inhibition of intravasation using a PI3K/mTOR inhibitor (proof-of-concept)

To validate the system as a drug-testing platform, we used Dactolisib, also known as BEZ235 or NVP-BEZ235, which is a dual inhibitor of PI3K (Phosphatidylinositol 3-kinase) and mTOR (mammalian target of rapamycin). The inhibitor was introduced via the perfusion medium and maintained in circulation for 24 hours under flow conditions. To evaluate the on-target effect of the inhibitor we performed immunofluorescence staining for phosphorylated S6 ribosomal protein (p-S6), a surrogate of PI3K/mTOR active signaling^21^. Confocal imaging showed the presence of p-S6 signal in tumor cells, which was reduced following inhibitor treatment (Figure 5A and B). Moreover, quantification of cancer cells revealed that Dactolisib reduced the intravasation score by five-fold (Figure 5C). In addition to efficacy, the drug safety assessment is a critical preclinical parameter during drug candidate development or validation. A key advantage of our system is the access to a functional endothelial lumen (vessel wall), which enables direct evaluation of vascular toxicity, a frequent dose-limiting side effect of anti-cancer therapies^22^. Quantification of the number of endothelial cells showed that Dactolisib at the concentration used (5µM), under shear stress, does not influence endothelial cell viability (Figure 5D).

**FIGURE 5.**
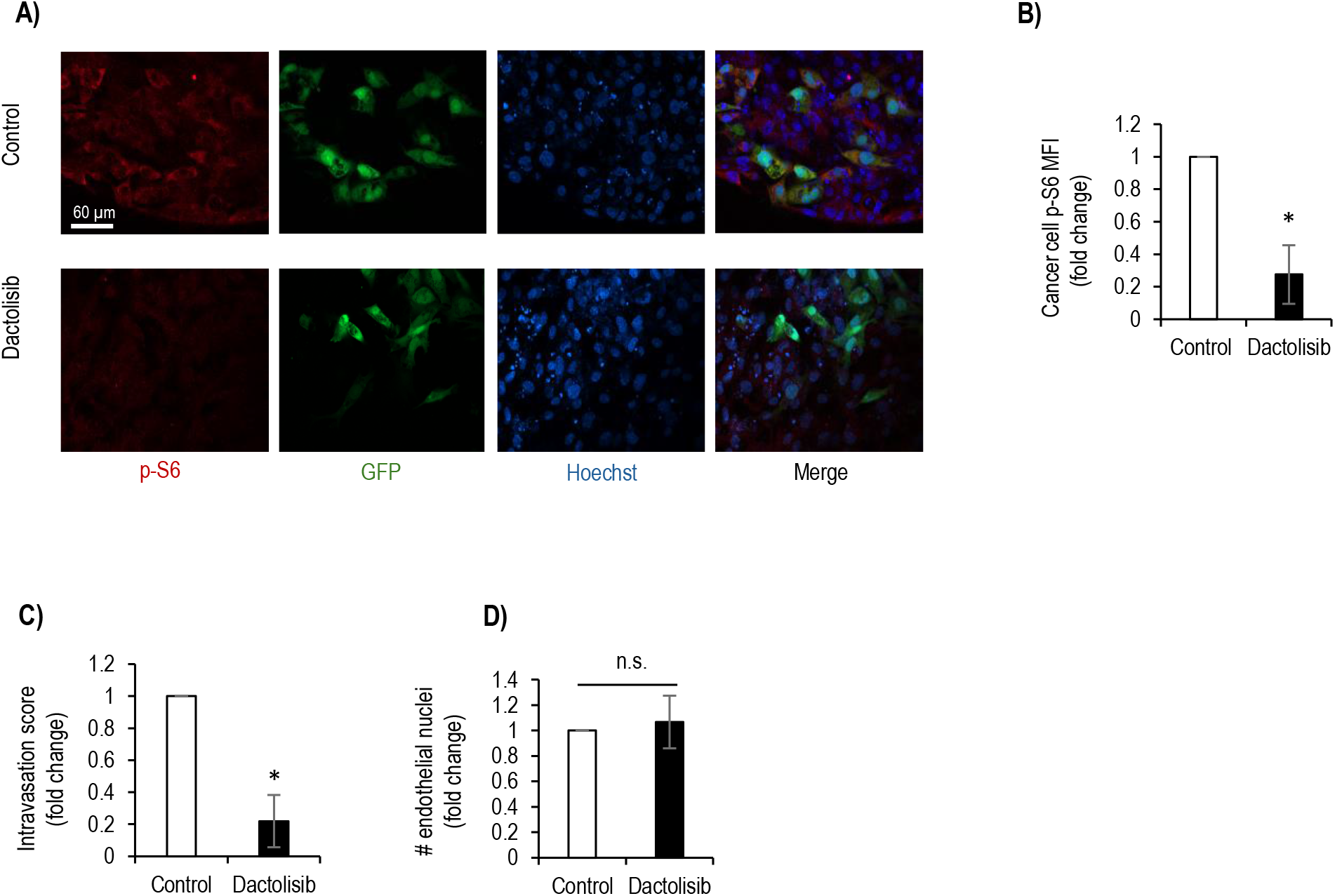
Intravasation inhibition – Proof-of-concept. **A)** Immunofluorescence staining of phosphorylated S6 (p-S6) in control and in the presence of Dactolisib. Scale bar = 60μm. B) Quantification of p-S6 in cancer cells in both conditions. The graph presents the mean ± standard deviation from three independent experiments. **C)** Intravasation scores in control and Dactolisib conditions. Intravasation scores were calculated by dividing the number of cancer cells present in the outlet plus the number of collected cells, by the number of cells in the inlet for each condition. **D)** Quantification of # of endothelial nuclei in both conditions. All graphs present the mean ± standard deviation from three independent experiments. Statistical significance was assessed by the two-sided t-test (* p<0.05).

## Discussion

Despite being one of the earlier steps of metastasis formation from solid tumors, intravasation is still a poorly understood biological process. A major reason for this is the scarcity of suitable *in vitro* models to mimic cancer cell transmigration through an endothelial monolayer to enter the blood circulation. Traditionally, transendothelial migration has been assayed in vitro using transwells or time-lapse imaging of transmigration with endothelial cells plated on top of tissue culture wells ^23^. These assays, however, better mimic extravasation rather than intravasation, as they assess the capacity of suspended cancer cells to adhere to the apical side of endothelial cells and transmigrate in an apical-to-basal direction. By contrast, during intravasation cancer cells are adherent and approach the endothelia from the basal side, transmigrating towards the vessel lumen. The development of vessel-on-a-chip allowed better modeling of this process by enabling cancer cells to be embedded in 3D matrices and migrate through endothelial barriers in a physiologically relevant basal to apical direction. Few intravasation-on-a-chip systems have been developed, and the dynamics of intravasation have been followed in real time with high resolution^20,24–26^. Moreover some models enable assessing intravasation under perfusion with variable speeds, mimicking the hemodynamic conditions that happen *in vivo*^17,20^. The five-channel configuration we developed allows chemoattraction studies putting in context the need for cancer cell directional migration prior to its intravasation, a key physiological aspect often ignored in other systems. By being sided by two ECM walls, cells have the possibility to choose to migrate to either side. In many platforms there is only one way cancer cells are able to invade the ECM ^24,26^. Our system features allow chemoattraction studies in the same chip: testing two experimental conditions simultaneously in the same device.

Another key advantage of the system is that it allows the quantification, isolation, and characterization of viable intravasated cancer cells. This is an important advantage, since similar setups only focus on the quantification of intravasation via *in situ* imaging methods of fixed or living samples ^16,20,24,26^. In the configuration we present, the one-way flow application allows the collection of viable intravasated cells at the end of the perfused vessel-like structure, opening the possibility for further downstream studies. The ability to sort cancer cells at distinct stages of invasion, intravasation, detachment from the endothelium, and circulation, followed by gene expression analysis, would enable characterization of transcriptional dynamics and provide insight into the molecular causes and consequences of vessel invasion by cancer cells.

Additionally, the system is well suited to identify anti-intravasation compounds with built-in capacity to simultaneously evaluate inhibitor on-target activity and endothelial toxicity, providing an integrated assessment of both therapeutic efficacy and vascular safety. As proof-of-concept, we used the PI3K/mTOR inhibitor Dactolisib. PI3K/mTOR signaling regulates multiple cellular processes including proliferation, migration, and invasion, in addition to transendothelial migration^21^. The outlet-to-inlet ratio metric accounts for these upstream effects, isolating the specific impact on migration and intravasation while correcting for possible effects of the inhibitor on proliferation. At the concentration tested (5µM) Dactolisib did not affect cancer cell proliferation (not shown); suggesting the reduced intravasation index primarily reflects decreased migratory or invasive capacity rather than reduced cell numbers.

There is a considerable need for more accurate human-representative systems to model the effects of drug candidate compounds on the body. Our platform enables simultaneous testing of anti-intravasation capacity, on-target effect, and vascular toxicity on the same chip. Overall, this system offers a powerful tool for mechanistic studies of intravasation and drug screening, with the potential to advance the development of anti-metastatic therapies. Future potential uses of the platform include the identification of the molecular mechanisms underlying intravasation.

Moreover, we identified refinements of the model that could enhance the physiological relevance and broaden the applications of this platform. The first, would be expanding the platform beyond the MDA-MB-231 triple-negative metastatic breast cancer model, to include other breast cancer subtypes (ER+, HER2+) and other cancers, to establish the platform’s versatility and confirm cancer (sub)type-universality of the metastatic process. Second, to replace HUVECs with organ-specific endothelial cells that more closely recapitulate the heterogeneity of the tumor vasculature encountered *in vivo*. Third, to evaluate alternative ECM compositions, since the reliance on Matrigel may be conditioned by batch-to-batch variability and the presence of xenogeneic components. Fourth, build on the model complexity by incorporating additional tumor microenvironment components, such as cancer-associated fibroblasts, immune cells (macrophages, T cells), and pericytes, would create a more complex and biomimetic system. This would leverage understanding to what extent the cellular crosstalk influences intravasation. Fifth, the modular assembly of this platform could be extended to model downstream metastatic events, for example, by connecting multiple organ-specific chambers, enabling the study of circulating tumor cell survival, extravasation, and organotropic colonization in an integrated multi-organ-on-chip system. Finally, the ability to test patient-derived cells directly in the chip could provide early patient stratification strategies.

## Methods

### Cell culture

Human umbilical vein cells (HUVECs) were obtained from Lonza (pooled), USA, and maintained in EGM-2 media (Lonza, USA) at 37 °C in 5% CO_2_ conditions. MDA-MB-231 and MCF-7-GFP expressing breast cancer cells were grown in Dulbecco’s Modified Eagle’s Medium (DMEM) supplemented with 10% FBS and 1% pen-strep under 37 °C and 5% CO_2_ conditions.

### Microfluidic chip design and fabrication

The five-compartment device, separated by small pillars, was designed using computer-aided design software to generate a photolithography mask. It comprised a central perfusable channel flanked by two lateral hydrogel chambers, which were, in turn, bordered by a pair of outer fluidic channels. It was fabricated using PDMS through replica molding from a SU-8/silicon master produced via ultraviolet photolithography. The PDMS base and curing agent were mixed at a 10:1 ratio, poured over the master mold, degassed under vacuum to eliminate air bubbles, and cured at 80°C for 2 hours. Following curing, the PDMS layers were cut using a razor blade, and fluidic ports (1.0 mm) were created using a biopsy punch. The PDMS devices were bonded to glass slides by oxygen plasma treatment (Harrick Plasma Cleaner, 1.75 Torr, 60 seconds) to ensure conformal contact and stable sealing.

### Migration and intravasation assay tumor-on-a-chip

Tumor cell seeding: Matrigel was injected into the second and fourth channels to serve as a supportive matrix and allowed to solidify at 37ºC. 1 × 10^5^ cells cancer cells were then seeded in Matrigel in the third channel of the microfluidic device using a micropipette. DMEM supplemented with 10% FBS was injected into the fifth channel, and Matrigel was added to the first channel. The device was covered with DMEM +10% FBS and incubated in a humidified atmosphere at 37 °C with 5% CO_2_.

### Endothelial cell seeding

Three days after cancer cell seeding, the fifth channel was coated with 2% Matrigel and incubated for 30 minutes at 37 °C with 5% CO_2_. HUVECs were then seeded into the same channel in four consecutive seedings. First, 0.25 × 10^5^ cells were seeded, and the chip was placed inverted at the incubator for 30 minutes to allow endothelial cell attachment to the top of the PDMS. Then, 0.25 × 10^5^ cells were seeded again, and the chip was tilted to the left side and kept in the incubator for 15 minutes. A third seeding was performed similarly on the opposite side, and finally, the chip was returned to its upright position, re-seeded, and incubated for an additional 30 minutes. At the end, the chip was covered with EGM-2 + 2% FBS media.

### Perfusion and CTC collection

Flow was initiated in the final right channel, where HUVECs had been seeded one day prior. EGM-2 medium was loaded into a syringe, and the tubing was connected to the inlets and outlets of the microfluidic chip. The outlet tubing was directed into a T-25 flask designated for collecting tumor cells, as shown in Figure 4A. Flow was applied in a stepwise manner: 1□L/minute for one hour, followed by 2□L/minute for two hours, 3□L/minute for three hours and finally 6□L/minute for 18 hours. Immediately after, the chips were fixed with 4% paraformaldehyde (PFA) in PBS containing calcium and magnesium at room temperature for 20 minutes.

### Drug testing

Dactolisib (at 5 μM in EGM-2) was perfused into the vessel channel for 24 hours following the previously detailed perfusion scheme. The chips were then fixed with 4% PFA diluted in PBS containing calcium and magnesium at room temperature for 20 minutes.

### Immunofluorescence staining

Immunofluorescence was performed by adding all the solutions to the vessel channel. First for permeabilization and blocking, a solution containing 3% bovine serum albumin (BSA) and 0.1% Triton X-100 in PBS was applied for 30 minutes. Subsequently, the channels were washed three times with PBS. Cells were then incubated overnight at 4 °C with the primary antibodies: anti-VE-cadherin (Santa Cruz, 6458) and anti-phosphorylated (Ser235/236) S6 Ribosomal Protein (Cell Signaling, 2211) prepared in a 1:100 dilution in 1% BSA in PBS. After incubation, the cells were washed three times with PBS to remove unbound antibody. Subsequently, they were incubated for 1 hour at room temperature with the appropriate secondary antibody (1:500 dilution) in 1% BSA in PBS. After the secondary antibody treatment, the cells were again washed three times with PBS. Nuclear staining was performed using Hoechst 33342, followed by a 15-minute incubation.

### Image acquisition and analysis

Live cell Imaging was performed in a Zeiss LSM 980 point-scanning confocal microscope equipped with a large cage and stage incubators (Pecon, Erbach, Germany) maintained at 37ºC and 5% CO2 with humidification. Images were acquired using an LD C-Apochromat 40x/1.1 water immersion objective. GFP fluorescence was detected using 488 nm for excitation and a 498-560 nm detection window. CellMask™ Deep Red fluorescence was detected using 639 nm for excitation and a 642-757 nm detection window. Z-stacks in a composition varying from 3 to 16 tiles of the two channels were acquired over a range of 109.92 µm. Image processing and 3D reconstruction were performed using Imaris software (version 9.9.1). This platform was used to generate 3D renderings and to acquire both still images and video recordings of the intravasation events.

Whole chip images (invasion assay) were acquired using a ZEISS Celldiscoverer 7 microscope. Images were captured using Plan-Apochromat 20x/0.95 objective for the MDA-MB-231 GFP^+^ chips and Plan-Apochromat 20x/0.7 objective for the MCF-7 GFP^+^ chip. The acquired images were stitched, and the preferential invasion quantification was performed by quantifying the GFP^+^ area fraction within the second and fourth channels of each chip using the ImageJ (2.16.0/1.54p) software. Images of immunofluorescence for VE-cadherin in endothelial cells were acquired in a Zeiss LSM 980 laser scanning confocal microscope at 40x magnification using LD LCI Plan-Apochromat 40x/1.2 water immersion objective.

For the quantification of cancer cells in the inlets and outlets Z-stack images in tiles were captured in a Zeiss LSM 980 laser scanning confocal microscope at 20 X magnification using Plan-Apochromat 20x/0.8 objective. The acquired images were stitched and processed and the quantification of MDA-MB-231 GFP^+^ cells was performed manually. For the analysis of phosphorylated S6 protein, one region of interest of the outlet was captured for each chip in a Zeiss LSM 980 point-scanning confocal microscope using Plan-Apochromat 20x/0.8 objective. Quantification of p-S6 in cancer cells was done using the ImageJ (2.16.0/1.54p) software by dividing mean intensity of p-S6 measured within a GFP mask by the number of GFP+ cells. All microscopy experiments were performed at the Bioimaging Platform of the Gulbenkian Institute for Molecular Medicine.

### Statistical analysis

The data were analyzed using two-tailed unpaired Student’s *t*-tests. Values are presented as mean ± standard deviation. Differences were considered significant when p<0.05 (*).

## Acknowledgments

We deeply thank all past and current members of the ADBCancer Lab for their dedication, enthusiasm and great team spirit. We are also grateful to Luís Costa’s Lab at GIMM for their invaluable technical and management support; Bruno Freitas and Afonso Malheiro for help with microfabrication, the João Barata Lab for reagents; and the technical support of GIMM’s Bioimaging Platform funded by PPBI-POCI-01-0145-FEDER-022122. This study was supported by Fundação para a Ciência e Tecnologia (FCT) of the Portuguese Government by I.P., under the project UIDP/50005/2020, by Fundo iMM-Laço and by GIMM-CARE. GIMM-CARE is funded by the European Union under grant agreement No. 101060102. GIMM-CARE is co-funded by the Portuguese Government, the National Foundation for Science and Technology (FCT), ARICA – Investimentos, Participações e Gestão, Jerónimo Martins, the Gulbenkian Institute for Molecular Medicine and CAML - Lisbon Academic Medical Centre. AM was also funded by Crédito Agrícola Vida – Companhia de Seguros, S.A. Personal research contract to K.S. (CEECIND/00697/2018) was supported by FCT.

## Author Contributions

Conceptualization: AM, SD, DB

Design of the experiments: AM, SD, DB, KS

Writing the paper: AM, NP, DC, CM, DB

Fabrication of the device: DB

Performing experiments: NP, DC, CM, AM, MF

Image acquisition and analysis: NP, DC, CM, MF

Lab management: DC, RN

Supervision: AM, SD, KS, DB, WR, EG, CF, LC

Discussion and revision: AM, SD, KS, DB, WR, EG, CF, LC, NP, DC, CM, MF, RN

Infrastructure: SD, KS, LC, EG, CF

Funding: AM, SD, KS

## Data availability

Data presented in this work is contained within figures, including supplementary and any clarification will be given upon request.

## Bibliography

1. Dillekås, H., Rogers, M. S. & Straume, O. Are 90% of deaths from cancer caused by metastases? Cancer Med. 8, 5574 (2019).

2. Sung, H. et al. Global Cancer Statistics 2020: GLOBOCAN Estimates of Incidence and Mortality Worldwide for 36 Cancers in 185 Countries. CA. Cancer J. Clin. 71, (2021).

3. Reymond, N., D’Água, B. B. & Ridley, A. J. Crossing the endothelial barrier during metastasis. Nature Reviews Cancer vol. 13 858–870 (2013).

4. Chiang, S. P. H., Cabrera, R. M. & Segall, J. E. Tumor cell intravasation. American Journal of Physiology - Cell Physiology (2016) doi:10.1152/ajpcell.00238.2015.

5. Sznurkowska, M. K. & Aceto, N. The gate to metastasis: key players in cancer cell intravasation. FEBS Journal (2022) doi:10.1111/febs.16046.

6. Kim, M. Y. et al. Tumor Self-Seeding by Circulating Cancer Cells. Cell 139, 1315–1326 (2009).

7. Zhang, W. et al. The bone microenvironment invigorates metastatic seeds for further dissemination. Cell 184, 2471-2486.e20 (2021).

8. Beinse, G. et al. Circulating tumor cell count and thrombosis in metastatic breast cancer. J. Thromb. Haemost. (2017) doi:10.1111/jth.13792.

9. Gi, T. et al. Histopathological Features of Cancer-Associated Venous Thromboembolism: Presence of Intrathrombus Cancer Cells and Prothrombotic Factors. Arterioscler. Thromb. Vasc. Biol. (2023) doi:10.1161/ATVBAHA.122.318463.

10. Boire, A. et al. Why do patients with cancer die? Nat. Rev. Cancer (2024) doi:10.1038/s41568-024-00708-4.

11. Newcomer, K. et al. Macrovascular tumor infiltration and circulating tumor cell cluster dynamics in patients with cancer approaching the end of life. Nat. Med. (2025) doi:10.1038/s41591-025-03966-3.

12. Sznurkowska, M. K. & Aceto, N. The gate to metastasis: key players in cancer cell intravasation. FEBS J. (2021) doi:10.1111/FEBS.16046.

13. Bockhorn, M., Jain, R. K. & Munn, L. L. Active versus passive mechanisms in metastasis: do cancer cells crawl into vessels, or are they pushed? Lancet Oncology (2007) doi:10.1016/S1470-2045(07)70140-7.

14. Sontheimer-Phelps, A., Hassell, B. A. & Ingber, D. E. Modelling cancer in microfluidic human organs-on-chips. Nat. Rev. Cancer 2019 192 19, 65–81 (2019).

15. Ingber, D. E. Human organs-on-chips for disease modelling, drug development and personalized medicine. Nature Reviews Genetics (2022) doi:10.1038/s41576-022-00466-9.

16. Shin, M. K., Kim, S. K. & Jung, H. Integration of intra- and extravasation in one cell-based microfluidic chip for the study of cancer metastasis. Lab Chip (2011) doi:10.1039/c1lc20671k.

17. Silvestri, V. L. et al. A Tissue-Engineered 3D Microvessel Model Reveals the Dynamics of Mosaic Vessel Formation in Breast Cancer. Cancer Res. (2020) doi:10.1158/0008-5472.can-19-1564.

18. Lee, H., Park, W., Ryu, H. & Jeon, N. L. A microfluidic platform for quantitative analysis of cancer angiogenesis and intravasation. Biomicrofluidics (2014) doi:10.1063/1.4894595.

19. Zervantonakis, I. K. et al. Three-dimensional microfluidic model for tumor cell intravasation and endothelial barrier function. Proc. Natl. Acad. Sci. U. S. A. (2012) doi:10.1073/pnas.1210182109.

20. Wong, A. D. & Searson, P. C. Live-cell imaging of invasion and intravasation in an artificial microvessel platform. Cancer Res. (2014) doi:10.1158/0008-5472.CAN-14-1042.

21. Glaviano, A. et al. PI3K/AKT/mTOR signaling transduction pathway and targeted therapies in cancer. Molecular Cancer (2023) doi:10.1186/s12943-023-01827-6.

22. Herrmann, J. Vascular toxic effects of cancer therapies. Nature Reviews Cardiology (2020) doi:10.1038/s41569-020-0347-2.

23. Reymond, N., D’Água, B. B. & Ridley, A. J. Crossing the endothelial barrier during metastasis. Nature Reviews Cancer (2013) doi:10.1038/nrc3628.

24. Zervantonakis, I. K. et al. Three-dimensional microfluidic model for tumor cell intravasation and endothelial barrier function. Proc. Natl. Acad. Sci. U. S. A. (2012) doi:10.1073/pnas.1210182109.

25. Lee, H., Park, W., Ryu, H. & Jeon, N. L. A microfluidic platform for quantitative analysis of cancer angiogenesis and intravasation. Biomicrofluidics (2014) doi:10.1063/1.4894595.

26. Ozer, L. Y., Fayed, H. S., Ericsson, J. & Al Haj Zen, A. Development of a cancer metastasis-on-chip assay for high throughput drug screening. Front. Oncol. (2023) doi:10.3389/fonc.2023.1269376.

